# Monkeys predict trajectories of virtual prey using basic variables from Newtonian physics

**DOI:** 10.1101/272260

**Authors:** Seng Bum M Yoo, Steven T. Piantadosi, Benjamin Y. Hayden

## Abstract

The demands of foraging are a major driver in the evolution of cognitive faculties. To successfully pursue a mobile prey that is attempting to avoid capture, the ability to predict its flight path can provide a crucial advantage. We hypothesized that, during pursuit, rhesus macaques exploit patterns in prey’s behavior to predict the prey’s future positions. We modeled behavior of three macaques in a joystick-controlled pursuit task in which prey follow simple escape algorithms that involve repulsion from the subject and from the walls of the virtual enclosure. We find that, even in this artificial task, macaques actively predict and aim towards prey’s future positions, increasing their foraging success. Their predictions are derived from the three core variables in Newtonian dynamics: position, velocity, and acceleration. Even after extensive training, subjects favored these principles and ignored other regularities in prey behavior. Most notably, they ignored the effects their own actions would have on the prey, despite extensive training and even though doing so would have further improved performance. We conjecture that subjects have a strong bias towards using physical principles to pursue fleeing prey, possibly reflecting an evolved physics module. The observed predictive behavior suggests that foraging demands facilitate the development of prospection.

## Significant Statement

Real-time prediction is crucial when chasing moving objects. We developed a novel virtual hunting task that requires macaque monkeys to control a joystick to pursue prey continuously moving on a computer screen. We found that subjects actively predict the upcoming position of the virtual prey by taking advantages of basic kinematic principles (speed and acceleration). Their predictions do not reflect expectations about the effects the agent’s own actions will have on the prey. These results demonstrate prospection in macaques and also suggest it may have practical limits.

## INTRODUCTION

The demands of foraging are a major driver of our neural and cognitive faculties, including specialized brain systems that allow us to perform complex computations in order to hunt more effectively (1, 2). When faced with mobile prey that move erratically, such as those that are fleeing, the ability to actively predict preys’ future positions can provides a boost in pursuit efficiency and, ultimately, survival (3–6). Prediction about the future movements of objects in the environment depends on the ability to calculate future events. It is therefore an element of a suite of skills that constitute prospection - a hallmark of human thought (7, 8).

In natural pursuit settings, the ability to predict—rather than simply follow—prey’s movement has been shown for some highly specialized species (9), but not primates. Nor are animal prospection abilities strongly established more generally (10–12). Well-known examples of putative animal prospection generally rely on naturalistic foraging contexts, suggesting that it is the need to forage that drives prospective abilities. Nonetheless, such tasks generally operate on the domain of long time scales (often, several days). We conjectured that prey-related prediction should be a much more general skill, and thus be readily observable at short time scales in rapidly changing environments.

The manner in which such prospection occurs is poorly understood. Sensorimotor control research shows that humans and animals use inverse models to generate the motor commands that are required to achieve desired sensory states and forward models to predict the sensory outcome of movement (13, 14). Those studies provide hints that prospection might require internal model that simulates predicted outcome. However, such models are limited to self-movement. The types of computations that we use to predict object movement remain unidentified. This problem is compounded when the predicted object adaptively avoids the subject and attempts to elude capture.

Here, we developed a virtual pursuit task to test rapid on-line prospection in macaque monkeys with a real-time adaptive pursuit component. The task is loosely inspired by the pursuit of insects, which are thought to be a major driver of primate evolution (15). To determine which strategy the subjects used to capture the prey, we developed a generative framework to model online pursuit movements. The model formalized different possible ways that subject might predict or follow prey, allowing us to quantitatively evaluate what determined subject movements. Our results suggest that macaque monkeys aim their joystick to the position of prey based on extrapolating physical variables of the trajectory, aiming at a position where physics predicts the object is going to be located, even in situations where prey’s motion has other regularities.

## RESULTS

We trained three rhesus macaque subjects on *computerized virtual pursuit task* (subjects K, H, and C). In our task, subjects used a joystick to control the position of an avatar (a yellow circle) moving continuously and smoothly in an open rectangular field on a computer screen (**Fig. 1** and **Methods**). On each trial, subjects had 20 seconds to pursue and capture a prey item (a fleeing colored square) to obtain a juice reward. Prey avoided the avatar with a deterministic strategy that combined repulsion from the subject’s current position with repulsion from the walls of the field (see **Methods** for details). One prey was present on each trial; the prey item on any trial was drawn randomly from a set of five, identified by color, that differed in maximum velocity and associated reward size (**Fig. 1**).

**Fig. 1.**
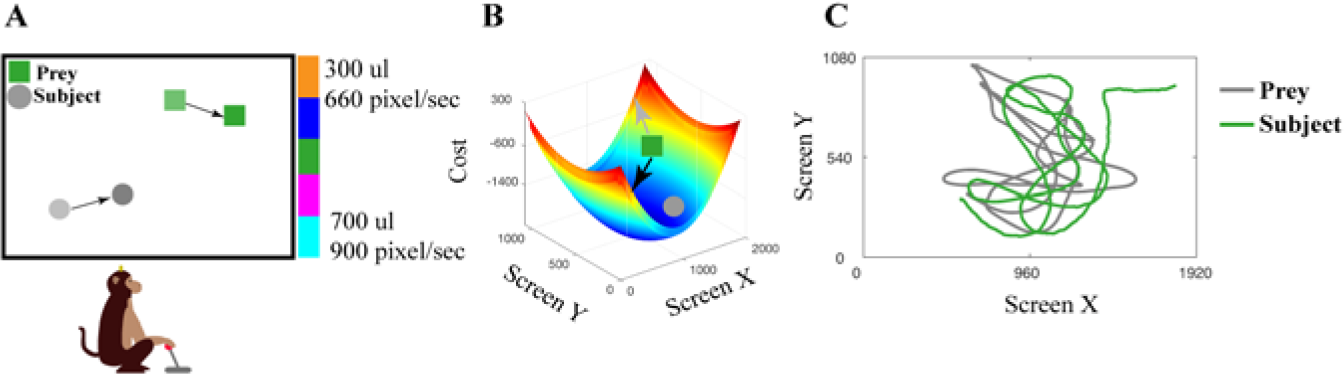
Experimental design. **(A)** Cartoon illustrating the virtual pursuit task. A subject moves a joystick to control the position of an avatar (yellow circle). The task code determines the prey’s next position according to the movement of the subject and the built-in cost contour map (in panel B). Updates of both subject and prey positions occur every 16.66 ms, (i.e. 60 Hz: identical to screen refresh rates). **(B)** The cost contour map across the screen. At each time-step, the task code determines the prey’s next move by choosing the position with the lowest cost. Cost is higher as prey moves closer to the edges and corners of the screen (to prevent the prey from being cornered by the predator). The configuration of this cost-contour map makes the prey more likely to move (1) away from the subject (grey arrow), and (2) towards the center of the screen (black arrow). **(C)** Example trajectories (capture time: 7.84 seconds). Green trajectory indicates the subject’s actual trajectory, while the grey trajectory indicates the prey’s actual trajectory.

All three subjects attained proficiency and showed stabilized behavior within twelve 2-hour training sessions that occurred following an initial longer training period on basic joystick use (see **Fig. S1** and **Fig. S2** for details). All data presented here were collected after the training sessions (number of trials analyzed in this post-training dataset: subject K: 3024; subject H: 3083; subject C: 2512). Subjects successfully captured the prey in over 95% of trials and, on successful trials, did so in an average of 5.04 seconds (subject K: 4.26 sec, subject H: 5.32 sec, subject C: 5.54 sec). Subjects’ performance and capture time declined with the maximum speed and reward offered by the prey, indicating sensitivity to manipulation of reward and difficulty (see **Fig. S1**).

We next fit the subjects’ strategies in pursuing their prey using a generative framework with a small number of parameters. Initially, the framework assumes that subjects exert a force towards either the prey’s future position (τ > 0), past position (τ < 0) or current position (τ = 0, **Fig. 2A**) using one of several possible predictive models of the prey’s motion. Our primary analyses thus determine (1) the value of the prospection parameter τ, and (2) the performance of different models of the prey’s predicted future motion, on each 1-second slice of the predator and prey’s motion. Along with τ, the strength of the force applied to the joystick, which in turn quantified the attraction to the prey, was a free parameter fit on each slice.

**Fig. 2.**
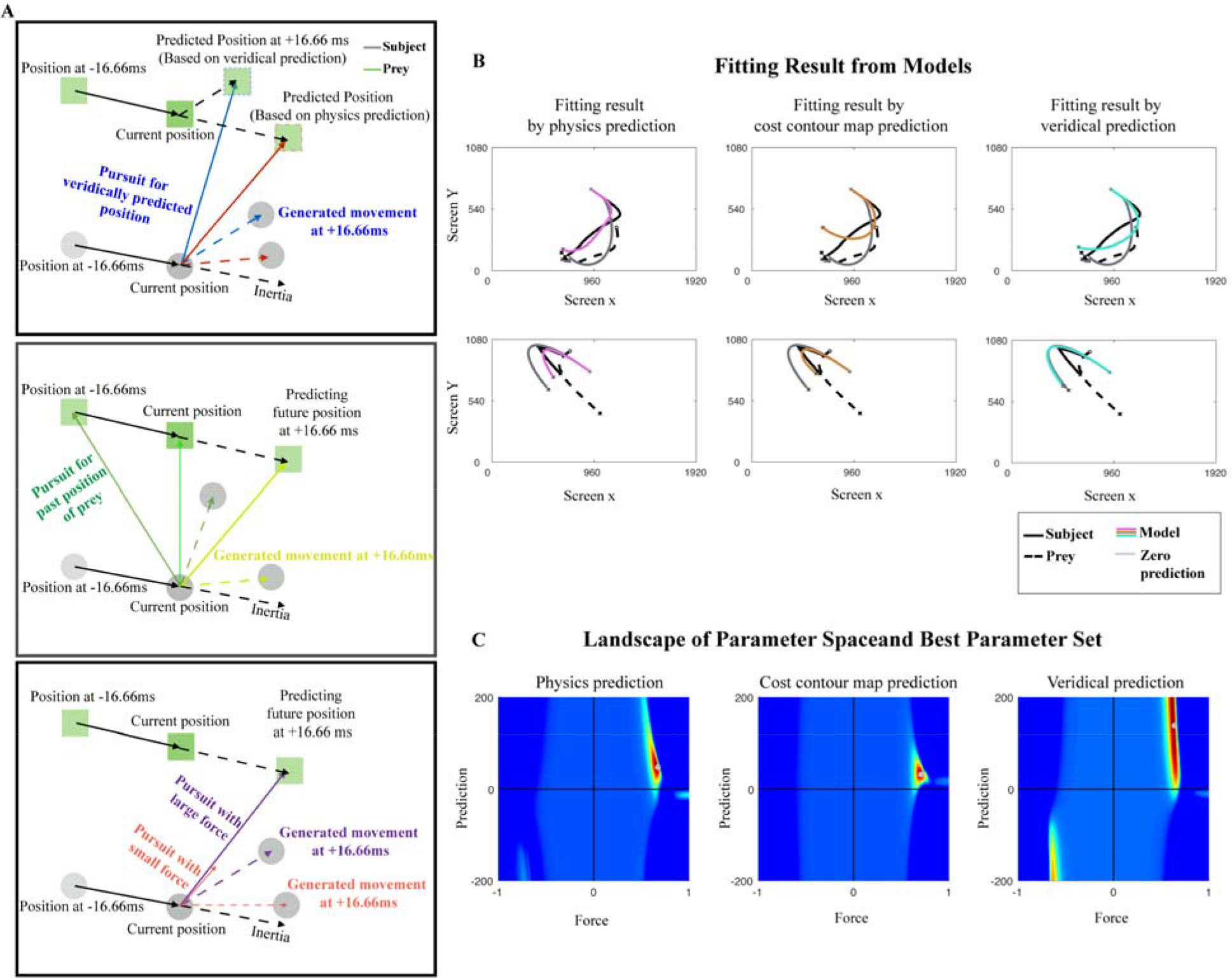
Model description, fitting results for single trial and population. (**A**) Model to generate trajectory based on prediction. Solid black arrow indicates movement from one previous time point (i.e. 16.66 ms before) to the current time point. The prediction for the prey’s position at the next time point is generated according to each model type. The monkeys are assumed to aim at a point forwards or backwards along the predicted trajectory, as determined by tau. The resulting movement vectors are constrained to a maximum speed specific to each subjects’ motor constraints. **(B)** Two example trajectories (left and right columns), and the fit trajectory generated by each prediction method (Physics Variable-Based, Cost Contour Map, and Veridical). The empty circle in the trajectory indicates the starting position, and the star marker indicates the ending position of the trajectory. **(C)** Heatmap plots of model performance explaining subject’s pursuit trajectory across parameter space from a single subject (Subject K). The gray circle indicates the best parameter combination explaining subject’s behavior, which generates the closest distance between the actual trajectory and model-predicted trajectory.

In addition to terms that define the amount of prospection and applied force, we added an inertia term reflecting physical constraints associated with our customized joystick. This inertia term was defined in terms of the velocity of the previous time step. The predicted movement is computed by summing a vector corresponding to the aimed direction and vector of inertia (**Fig. 2**). To compare the difference in fitting result, we calculated the difference of ‘the mean for sum-of-squared error’ between the model in each trajectory. Thus, if the model with inertia explained specific trajectory better, the value should be negative. We found that in all cases, subjects’ performance was better explained by the model having inertia term (**Fig. 3**). For significance testing, we performed bootstrapping for the value we obtained. We still found 5% significance line resulted in a negative value, meaning the negative value obtained in here is significant.

**Fig. 3.**
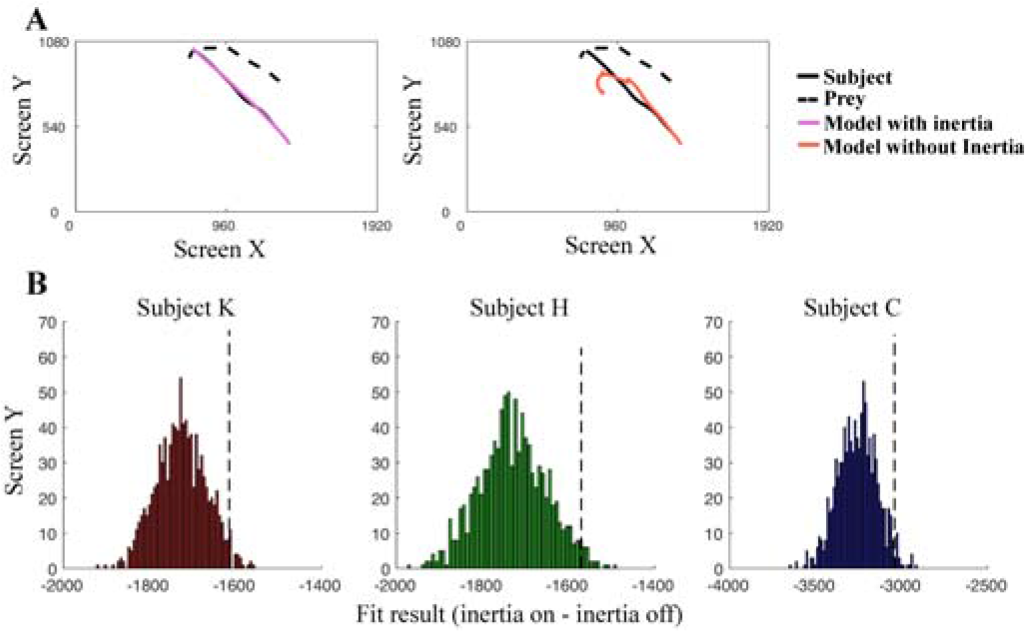
Inertia enhances model performance. (**A**) Model trajectory comparison between models with and without inertia. (**B**) Histogram results suggest that incorporating inertia component to the model leads to a better fit of the data (mean for sum-of-squared error difference below zero at x-axis). 95% of data falls to the left of the black, dashed line. Bootstrapping of difference in performance between the model with and without inertia was performed in randomly sampled trajectories (number of resamples: 1000, randomly selected trajectories: 2000).

To quantify the typical parameter values, we averaged a full grid of parameter values across trajectories, shown in **Fig. 2C**, using a *physics-variable based prediction model* (PVBP) of prey motion (see **Methods**). We observed a strong preference for positive force, demonstrating monkeys are engaging the task. The best fitting τ is positive, indicating that subjects point the joystick towards the prey’s future trajectory. This pattern holds for all three individuals tested. Specifically, subjects K, H and C pointed the joystick towards the position at which the prey would be in an average of 800 ms, 767 ms, and 733 ms, in the future respectively. All of these times are significantly greater than zero (more than 95% of bootstrapped data stayed above zero). In the context of the task, these numbers are substantial: they reflect 18.78%, 14.42%, and 13.23% of the average trial duration for K, H, and C, respectively.

The distance into the future that our subjects prospected did not reliably depend on the reward or the speed of the prey, as measured using a linear regression between reward/speed and mean τ (K: r = 3.0316, p = 0.1110; H: r = 4.5798, p = 0.1791; C: r = 7.1007, p = 0.0957). Prey path complexity (as measured by path curvature) did affect prediction. Subjects prospected less far into the future when the prey path was more complex (K: rho = -0.0687; H: -0.0567; C: -0.0898, p < 0.0001 for each).

We next quantitatively compared possible strategies subjects used to predict future prey direction by formalizing different computations by which monkeys could predict future trajectories (**Fig. 2A**) and fitting the parameters to each. The *veridical prediction (VP) algorithm* assumes that monkeys predict according to the true game dynamics in which prey move away from the boundaries of the field and also from the avatar. The *cost contour map prediction (CCMP) algorithm* matches VP but ignores repulsion from the avatar, meaning that monkey’s model of prey would not take into account their own motion. Third, the *physics variable-based prediction (PVBP) algorithm* assumes that subjects’ predictions derive from the prey’s position and first two derivatives, velocity and acceleration (additional derivatives are considered in **Fig. S3**). We measured the accuracy of each algorithm by computing the predicted path of the subject on every trajectory slice then computing its error (sum of squared distance between predicted and observed trajectories).

We use the Akaike Information Criterion (AIC) to compare models (**Fig. 4** and **Methods**). This figure shows that the PVBP model of future prey trajectories is overall the best fit to our subjects’ behavior. This pattern held within the two well-trained subjects. Specifically, the *PVBP algorithm* was favored (Subject K: PVBP: 7.529×10^6^, second best was VP: 7.542×10^6^; Subject H: PVBP: 8.923×10^6^; second best was CMPP: 8.950×10^6^, **Fig. 4A**). For the less well-trained subject C, *CCMP* explained trajectories most accurately (7.955×10^6^, VP: 8.013×10^6^). These patterns appear to be robust to the specific analysis as they could be seen in also by estimating which model fit best for each individual trajectory slice (**Fig. 4B**).

**Fig. 4.**
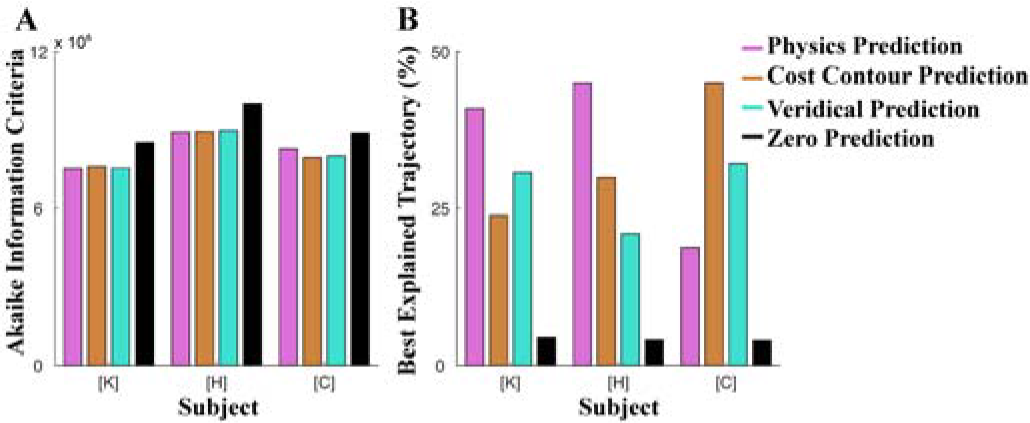
The fitting result across the subjects and model comparison. (A) Normalized Akaike Information Criteria (AIC) across all the trajectories. In the current figure, only the Physics Variable-Based Prediction (PVBP) based on velocity and acceleration is included as representative of all the physics variable-based prediction models. (B) Percentage of trials best explained by each model. The AIC values are compared across models.

Because Subject C showed a different pattern, and Subject C performed worst overall (and thus chased slower prey), we wondered whether prey speed may influence strategy. Supporting this idea, a trial-by-trial logistic regression between whether PVBP was the best model and average prey velocity showed a positive relationship for all three subjects (p < 0.01 in each case), with subject C maintaining a similar proportion of trajectories best explained by PVBP for its speed (**Fig. S5**). These results highlight the adaptive flexibility of prospective pursuit strategy selection, and indicate that Subject C’s overall difference can be explained by the relatively slower speed prey used.

We asked which values of parameters are closest to optimal in capturing prey using simulations (**Fig. 5**). To exclude the possibility where optimal parameters exist beyond what subjects can accomplish using the current joystick configuration, we examined optimality by comparing performance under identical pursuit/inertia ratios, which can be accomplished by limiting the range of the force parameter in simulation. The representative prediction parameter in simulation shows that all subjects’ prediction parameter sets are not identical to the optimal parameter set obtained from PVBP simulation. Average capture time in simulation using optimal parameter was 1.10 second while top 5% capture time of actual trial was 1.31 second (subject K), 1.32 second (subject H). The value of the optimal prediction parameter was 335 pixels compare to actual prediction of each animal was 536 (subject K), 455 (subject H) pixels (Subject C’s prediction is not directly comparable in pixel units because the prey in simulation had maximum speed matched with subject K and subject H but faster than subject C’s game). These results suggest that our subjects’ pursuit strategy is less than the optimal even subject to reasonable empirically derived constraints, and performance would have improved once they used shorter prediction scales.

**Fig. 5.**
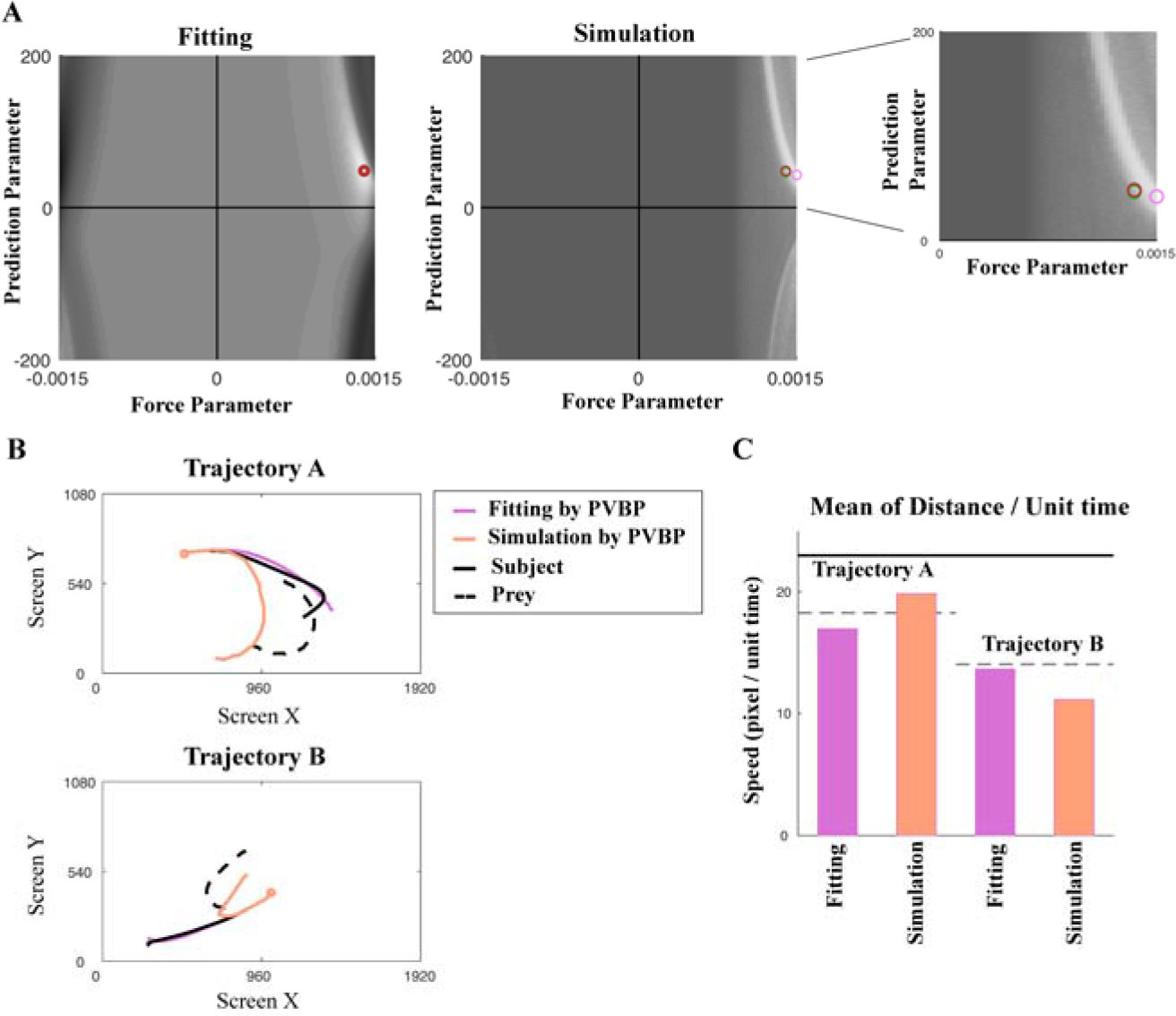
Pursuit trajectory of monkey is less than optimal compare to simulation result. (**A**) Left column is heatmap of fitting trajectory (identical as Fig. 2C, gray scaled for showing subject’s best parameter in color) and right column is capture time result from simulating artificial subject. Pink circle indicates best parameter obtained from simulation and other circles is best parameter from fitting each subject (red: Subject K; green: Subject H; subject C was not included since simulation matched prey speed with Subject K and H). (**B**) Trajectories generated by using best parameter from fitting (pink) and best parameter from simulation (light orange). The results suggest different strategy selection according to parameter (Quantitative comparison for capturing time between simulation and actual monkey’s behavior is at supplementary results). (**C**) Mean of distance per unit time in given example trajectory. Black line is maximum distance per unit time possible and gray dashed line is the mean of actual subject’s distance per time. Fitting parameter results in smaller value for trajectory A while has larger value for trajectory B.

## DISCUSSION

We trained macaques to perform a novel joystick-based pursuit task. The generative framework we developed to model our subjects’ strategies shows that macaque monkeys do not simply point the joystick towards the prey, but actively predict its future position. Notably, they extended basic principles of Newtonian physics to this artificial situation. That is, each subject’s movement is best explained by a model that predicts prey position based on extrapolating variables derived from physics (velocity and acceleration). These results therefore show that, in a difficult dynamic pursuit context, macaques will rapidly and naturally take advantage of regularities in their prey’s behavior to gain an advantage in reward intake. Moreover, they show that our subjects demonstrate a basic ability to perform the necessary computations to predict future prey positions and to exploit that prediction.

Our generative framework does not explain why animals prefer using basic physical principles and ignoring other factors that could make their predictions more accurate. One possibility is that the brain is equipped with a physics module that performs mental simulation about the physics of external environment (16). By having a module that specifically simulates physics, computations for external physics change would be more efficient than reasoning about underlying method how prey movement is generated. The fact that animals favor the physics predictions in the most difficult cases – those with the fastest prey – provides at least suggestive evidence for the idea that the physics based approach is one that is more natural.

Our results provide support, in a very different form than other studies, for the idea that our brains simulate physics to making judgement of the scene, sometimes called ‘intuitive physics’ (17–20). Previous studies related to intuitive physics have mainly focused scene understanding or moving objects without any interactions between the subject participant and the stimulus. Our study expands on these previous findings by generalizing for cases where environment changes dynamically and interactively.

More broadly, our results open a unique direction to understand real-time pursuit behavior. The majority of the behavioral studies try to use minimally simple tasks that sacrifice naturalness in order to control each variable precisely to understand behaviors. Such tasks can cause behavior to enter into modes that are unnatural. On the other hand, real-time naturalistic behavior requires frameworks to narrow down hypothesis space. Our study has set example of studying real-time dynamic behavior under guidance of well-established framework.

Questions about prospection aside, the ability to make choices based on expectations of future events is a basic skill in the repertoire of intelligent decision-makers (21–23). Those abilities guide appropriate selection of choices in foraging contexts, including under both risk and delay (24–26). A unique factor of our study is its focus on real-time decisions, that is, decisions in which subjects choose from a continuum of possible actions - which constitute a corresponding continuum of options - and reassess their options as their actions occur (27–30). Some scholars have argued that these types of decisions are the type of decisions that drove the evolution of our choice systems (2, 31, 32). As such they provide a more realistic assessment of choice behavior than binary choice tasks. We anticipate that future studies using this paradigm and others like it will lead to greater insight into the psychological and neural mechanisms of choice.

## MATERIALS AND METHODS

### Subjects

Three male rhesus macaques (Macaca mulatta) served as subjects in the current experiment. All animal procedures were approved by the University Committee on Animal Resources at the University of Rochester and were designed and conducted in compliance with the Public Health Service’s Guide for the Care and Use of Animals.

### Experimental Apparatus

The joystick was a modified version of commercially available joysticks with a built-in potentiometer (Logitech Extreme Pro 3D). The control bar was removed and replaced with a control stick (15 cm, plastic) designed to be ergonomically easy for macaques to manipulate. The joystick position was read out by a custom coded program in Matlab running on the stimulus-control computer. The joystick was controlled by detecting the positional change of the joystick and limiting the maximum pixel movement to within 23 pixels in 16.66 ms.

### Task Design

At the beginning of each trial, two shapes appeared on a gray computer monitor placed directly in front of the macaque subject. The yellow (subject K) and purple circle (subject H and C) shape (15-pixel diameter) represents the subject himself and its position is determined by the joystick and limited by the screen boundaries. The square shape (30-pixel length) represents the prey. The movement of the prey is determined by a simple AI (see below). Each trial ends with either successful capture of the prey or after 20 seconds, whichever comes first. Successful capture is defined as any overlap between the avatar circle and the prey square. Capture results in immediate juice reward; juice amount corresponds to prey color: orange (0.3 mL), blue (0.4 mL), green (0.5 mL), violet (0.6 mL), and cyan (0.7 mL).

The path of the prey was computed interactively using A-star pathfinding methods, which are commonly used in video gaming (33). For every frame (16.66 ms), we computed the cost of 15 possible future positions the prey could move to in the next time-step. These 15 positions were spaced equally on the circumference of a circle centered on the prey’s current position, with radius equal to the maximum distance the prey could travel within one time-step. The cost in turn is computed based on two factors: the position in the field and the position of the subject’s avatar. The field that the prey moves in has a built-in bias for cost, which makes the prey more likely to move towards the center (**Fig. 1B**). The cost due to distance from the subject’s avatar is transformed using a sigmoidal function: the cost becomes zero beyond a certain distance so that the prey does not move, and the cost becomes greater as distance from the subject’s avatar decreases. Eventually, the costs from these 15 positions are calculated and the position with the lowest cost is selected for the next movement. If the next movement is beyond the screen range (1920×1080 resolution), then the position with the second lowest cost is selected, and so on.

The maximum speed of the subject was 23 pixels per frame (i.e. 16.66 ms). The maximum and minimum speeds of the prey varied across subjects and were set by the experimenter to obtain a large number of trials (**Fig. 1**). Specifically, speeds were selected so that subjects could capture prey on above 85% of trials; these values were modified using a staircase method. If subjects missed the prey three times consecutively, then the speed of the prey was reduced. Once the subject intercepts the prey in a trial where the staircase method was used, then the selection of prey speed was randomized again. To ensure sufficient time of pursuit, the minimum distance between the initial position of each subject avatar and prey was 400 pixels.

### Behavioral Model

To fit each subject’s movement, each trial was divided into 1 second-long trajectories and each trajectory included 61 data-points with 16.66 ms time resolution. We model these trajectories using a single prediction and a single force parameter for the entire trial, as a simplifying assumption. Nonetheless, it is reasonable to assume that throughout a long, 20-second period, there would be active adjustment of prediction and force. Actual comparison by AIC supported our intuition, and we used trajectory as unit of analysis throughout (value of ‘AIC of trajectory/AIC of trial’ was 0.9328, 0.9214, 0.9227, for each subject respectively. Less than one indicates trajectory as unit result in better fitting).

The model assuming Physics Variable-Based Prediction (PVBP) incorporated one previous time step to predict the prey’s next position, which is like a Kalman filter. The other two models do not utilize any past information. The model assuming prediction using the cost contour map considers only the lowest cost location at next time step. The model assuming veridical prediction automatically finds the exact position of the prey at the next time step. Once the prey’s position on the next time step is predicted, the model computes how far this predicted position is from the agent’s current position. A prediction value of 1 indicates that the future position will be as far as from the agent’s current position as the prey’s current position is. The optimal parameter pairs of how much subject has made prediction and actual amount of force was acquired by performing a grid search across the ranges of both parameters. The range of the prediction parameter was between -400 to 400 subjects H and C, -200 to 200 for subject K (units were defined relative to the distance the prey moved in the previous timestep). Different ranges of the prediction parameter were used since over 5% of trajectories in subjects H and C resulted either in -200 or 200 in prediction parameter value. Representative parameters for explaining an each trajectory were selected based on the value of the sum of squared error between the actual trajectory and the trajectory generated by model.

### Significance Testing

To see whether the positive prediction parameter is significant above the zero, we performed a bootstrap of heatmap slices from each trajectory. This resampling was done for 500 times and selected heatmaps were added. Then, the parameter set resulting in the lowest cost was selected in each resampling.

### Model Evaluation

To evaluate model performance and compare different models, we computed the Akaike Information Criteria (AIC) using the likelihood of each model (**Fig. 4, and Fig. S3, Fig. S4**). For calculating likelihood, we first calculated the mean and variance of all the sum-of-squared errors across trajectories. Then we estimated the likelihood assuming a normal distribution centered on the mean of the sum-of-squared errors and with a variance equivalent to the variance of the sum-of-squared errors across all trajectories. To validate whether subjects used a single prediction and force across the all the trials or adaptively changed their prediction method, we compared the AIC value between cases where the parameter pair varied across all trajectories and using only the single best parameter pair. The single best parameter pair is acquired by lowest cost value in all trajectory added heatmap of parameter space.

### Simulation

To estimate the efficiency of parameter values obtained from fitting subjects’ behavior, we performed a simulation with 100 different initial positions of artificial subjects and prey (**Fig.5**). The algorithm generating prey movements was the same as that used in the actual task, and the movements of the artificial subject were generated based on different prediction methods. The maximum duration for each simulation was 1200 time-bins, which is equivalent to 20 seconds (the longest possible trial in the actual task). All the values of the parameter sets (prediction, force) that were used for fitting actual behavior were simulated.

### Velocity Dependent Physics Variable-Based Prediction Bias

We examined whether PVBP is preferred when the velocity of prey is high. We first obtain the average velocity of prey at each trajectory, and then categorized each trajectory as physics variable-based prediction if the fitting result was best with physics variable-based prediction and non-physics when other prediction method provided the best fitting result. With the prey velocity and trajectory category, we performed logistic regression having velocity as predictor and category as the dependent variable (glmfit in MATLAB).

### Data availability

The data sets generated during the current study are available on the Hayden lab website, http://www.haydenlab.com/, or from the authors on reasonable request. The code generated to do the analyses for the current study is available from the corresponding author on reasonable request. Video of experiment is available at http://www.haydenlab.com/pursuit.

## Acknowledgements

We thank Habiba Azab, Steve Chang, and John Pearson for the valuable discussions and comments in manuscripts. We are grateful for Haydenlab and CoLaLa lab for intuitive discussion that has lead into elaborating the model classes. We appreciate Marc Mancerella and Giuliana LoConte for advising the training monkeys, Alex Thome for developing method, and Shannon Cahalan for helping data collection. This work was supported by the NIH R01 DA038615 for B.H.Y.

## Authors Contribution

S.B.M.Y. and B.Y.H. designed the study. S.B.M.Y. trained monkeys and collected data. S.P. and S.B.M.Y develop the behavioral model, analyzed the data and model outcome. B.Y.H., S.P. and S.B.M.Y. wrote the manuscript.

## Competing interests statement

The authors declare that they have no competing financial interests.

**Correspondence** and request for the materials should be address to S.B.M.Y

## SUPPLEMENTARY MATERIALS

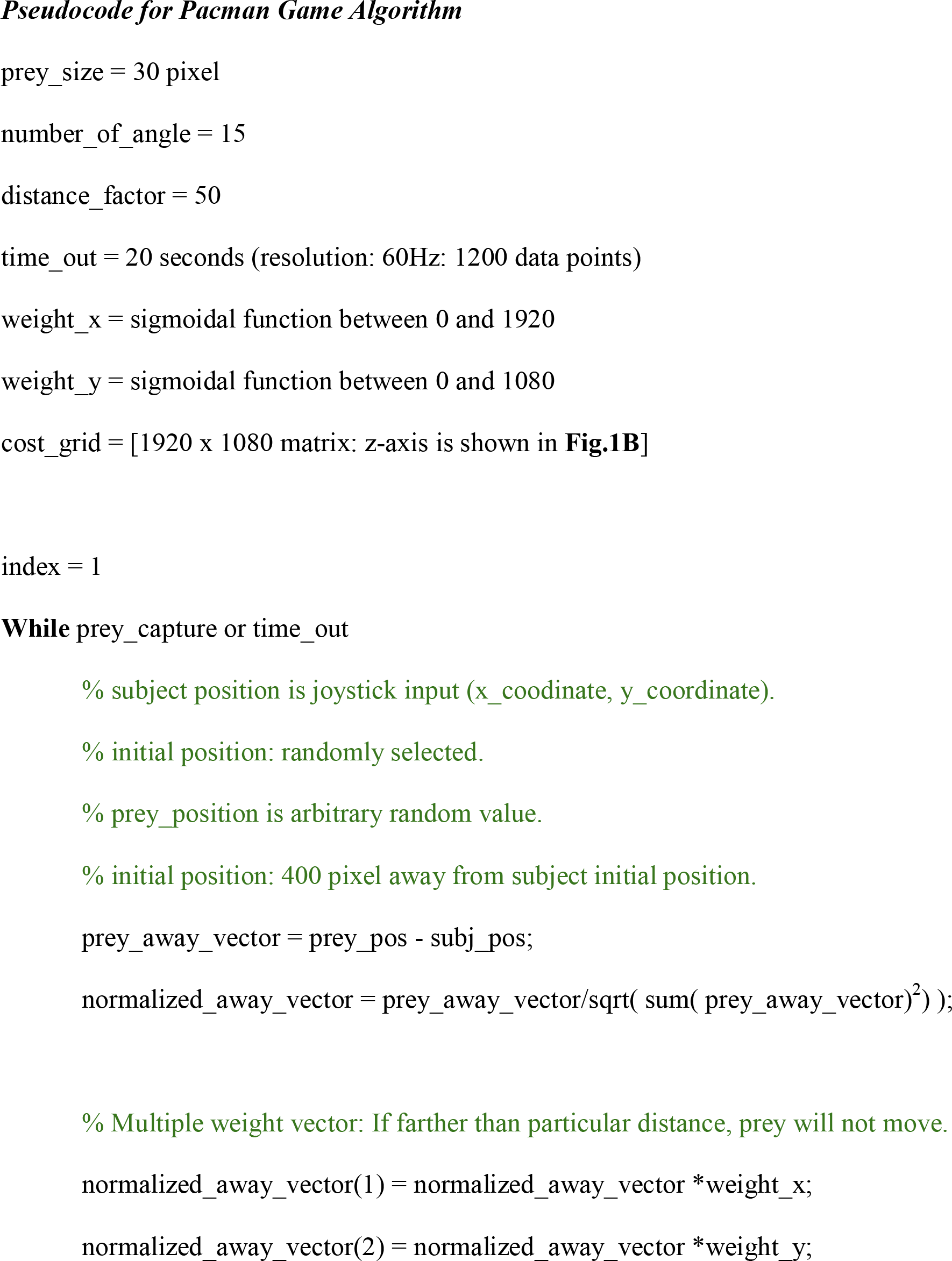

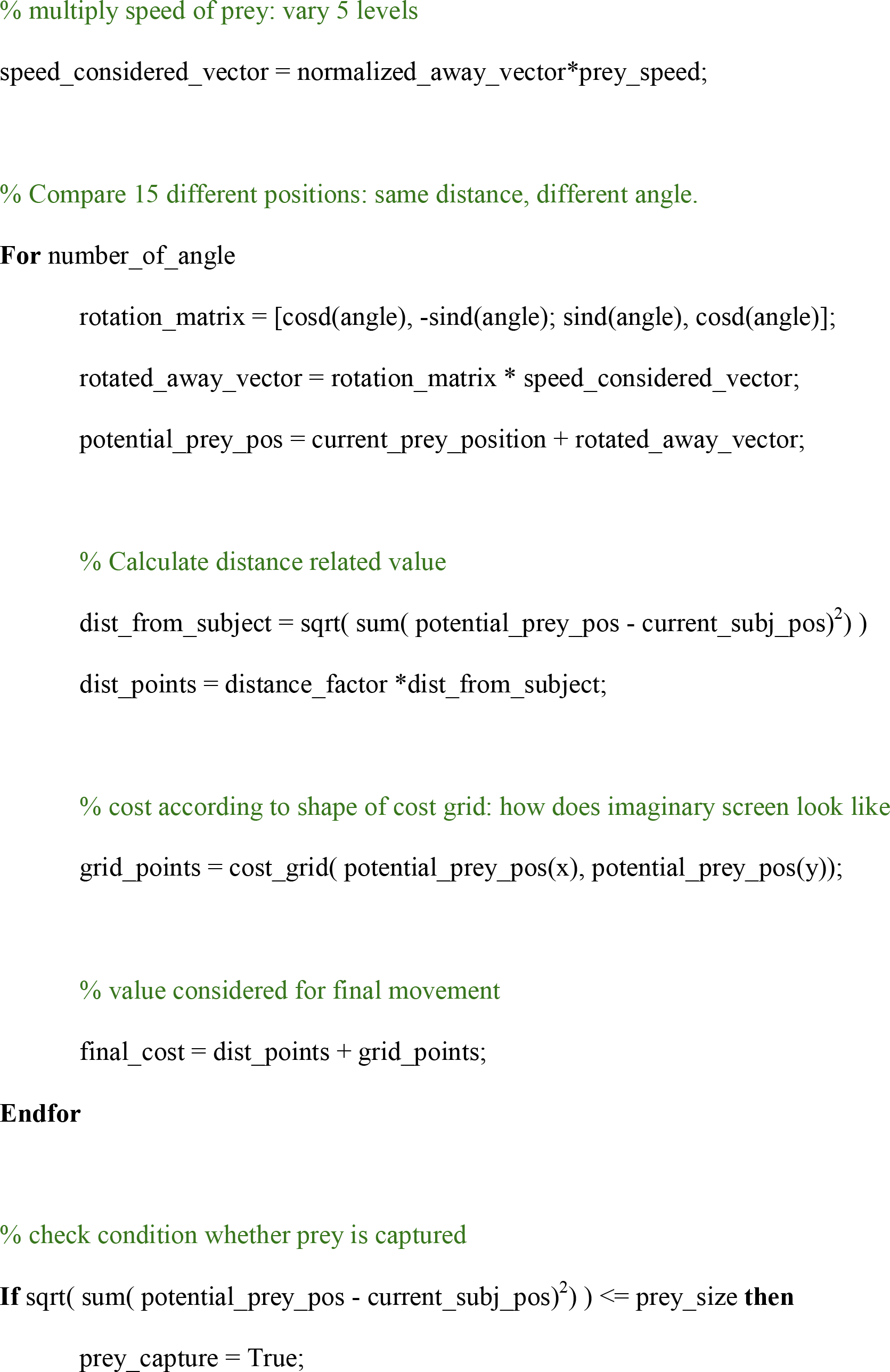

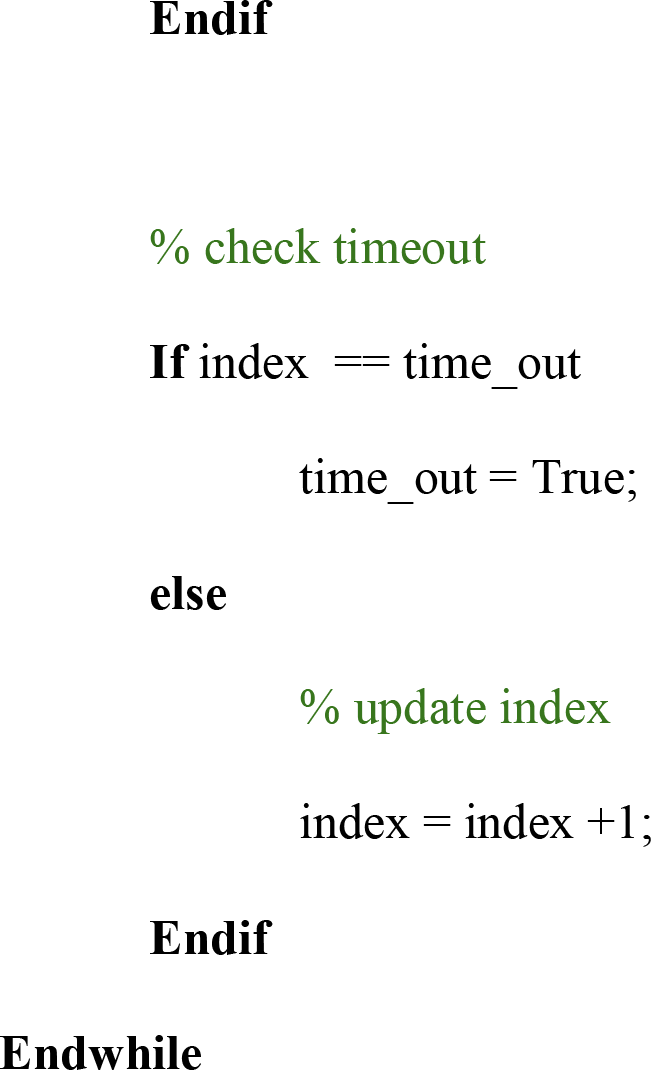

**Fig S1.**
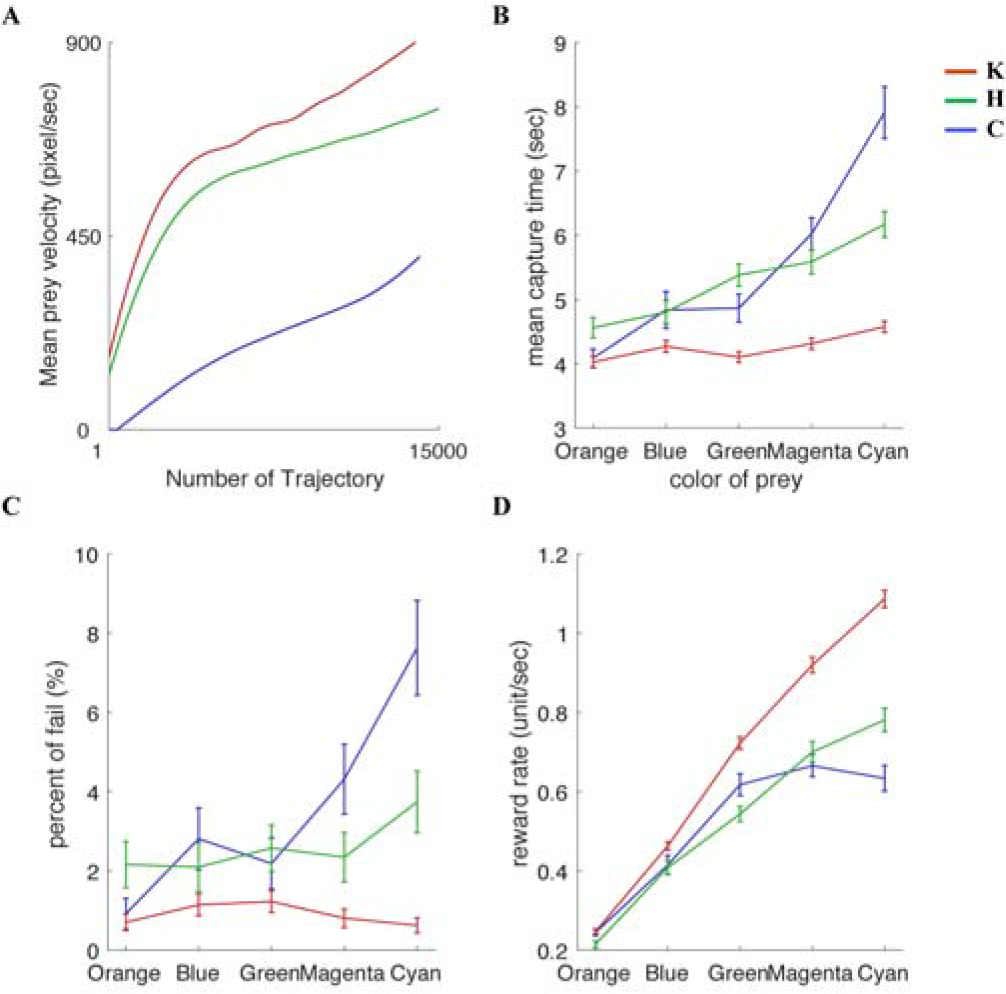
Monkeys behavior varies according to prey speed/reward. (**A**) Mean prey velocity in each trajectory plotted separately for each subject. Pursuit result differs according to color (equivalent to maximum speed) of prey. The maximum speed of prey increases from orange (slowest with smallest reward) to cyan (fastest with largest reward). As maximum speed increases, the mean capturing time (**B**) and percent of failed trials increase (**C**). However, reward rate also increases since the amount of reward is larger for faster prey (**D**). Errorbars are the standard error of the mean, obtained by bootstrapping (1000 bootstraps).

**Fig S2.**
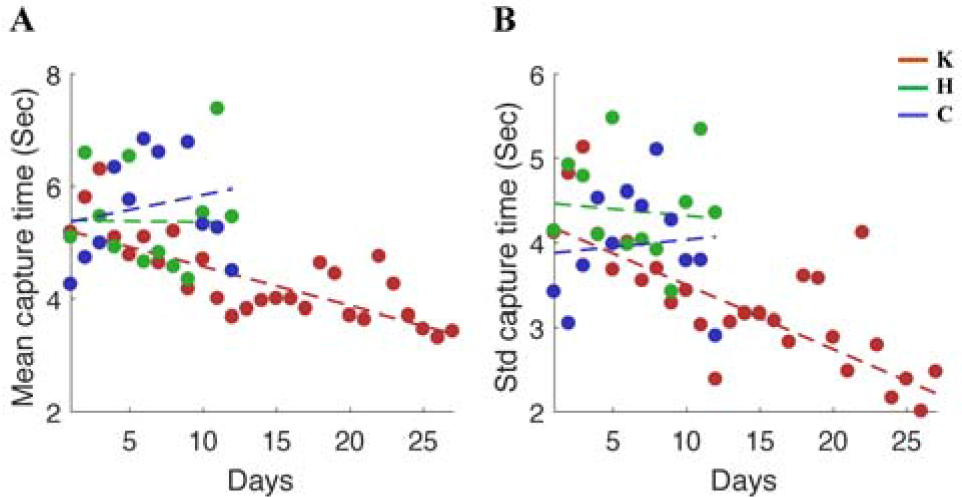
Stabilization of the behavioral performance. Performance stabilization shown by the mean (**A**) and the standard deviation (**B**) of prey capture time. Subject C (blue) and Subject H (green) do not show any significant changes in their capture time throughout the days of the experiment (linear regression with p-value > 0.1) while Subject K (red) shows significant changes to both mean and standard deviation of capture time across days (p-value < 0.001).

**Fig S3.**
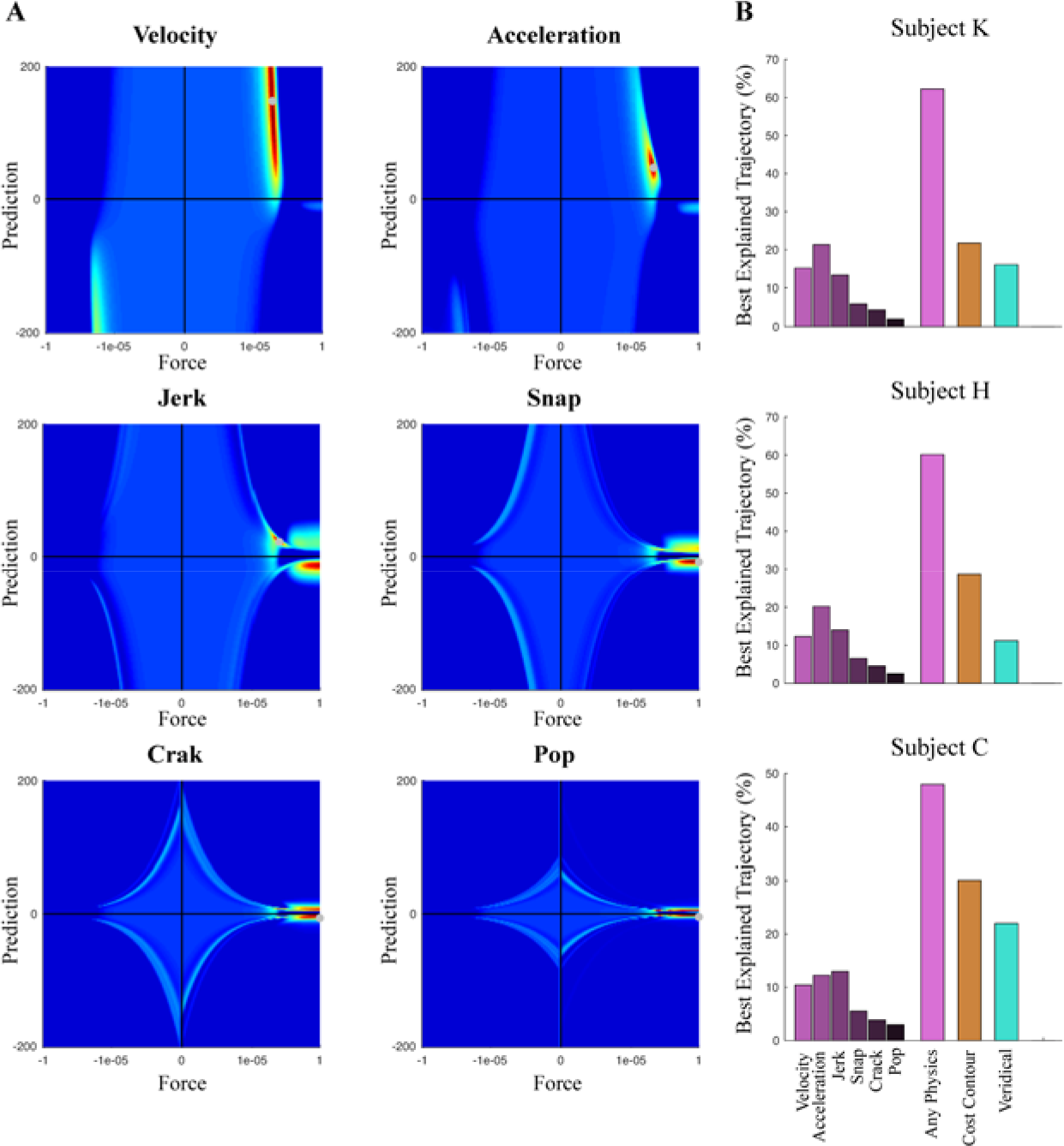
Second order approximation of physics explains trajectory most accurately. (**A**) Each heatmap indicates the addition of more physical derivatives until sixth derivatives. The black circle indicates the best parameter set for the model. (B) In summary figure, physics include within-physics prediction model comparison (from velocity to pop, the 6th derivative). Any physics indicate summed result of whole physics variable based prediction (PVBP) model class to compare with other prediction methods.

**Fig S4.**
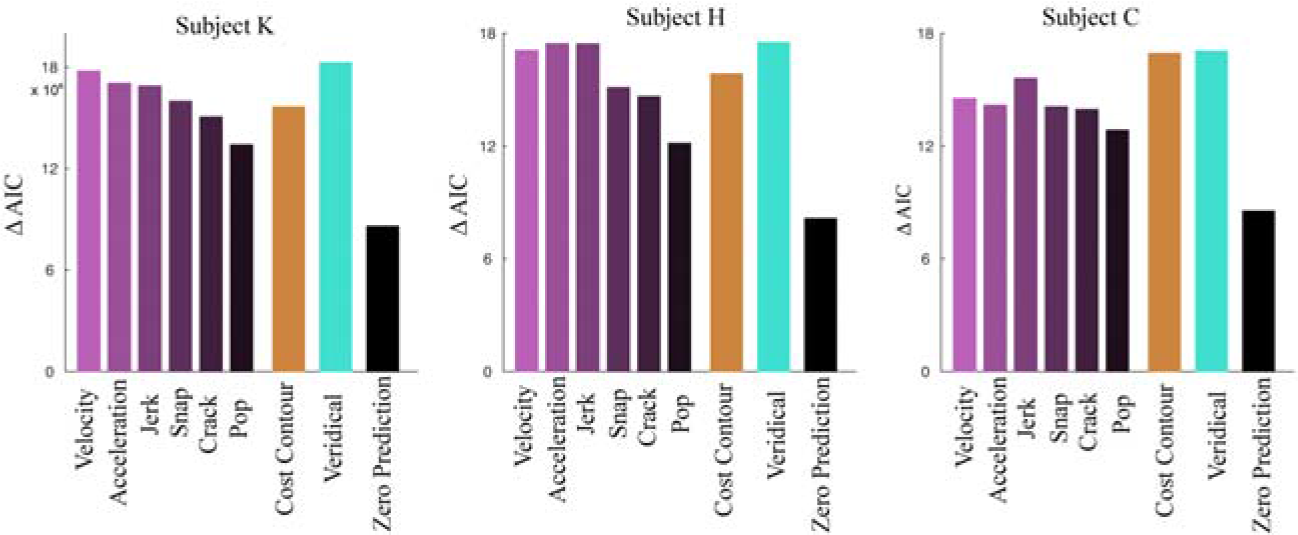
Dynamic change of parameter set at each trajectory explains monkey’s trajectory better than identical single parameter set across all the trajectories. AIC comparison between the case of the single parameter set across all the session (case 1) or adaptively changing parameter set at each trajectory (case 2). Delta AIC indicates the difference between the cases (case 1 - case 2), and a positive value indicates adaptively changing the strategy explains subject’s trajectory better even there is penalty having more parameters. Each column shows the individual subject result.

**Fig S5.**
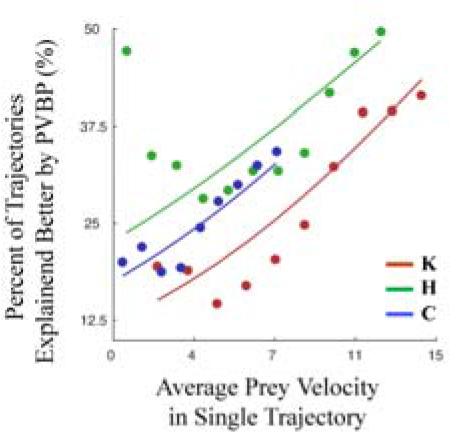
Prey velocity dependent strategy selection. All the monkeys consistently show biases using PVBP when the prey velocity is faster. Logistic regression was performed between prey velocity and categorical dependent variable (0: non-PVBP, 1: PVBP). The p-values of all logistic coefficient was significant (p < 0.001).

**Fig S6.**
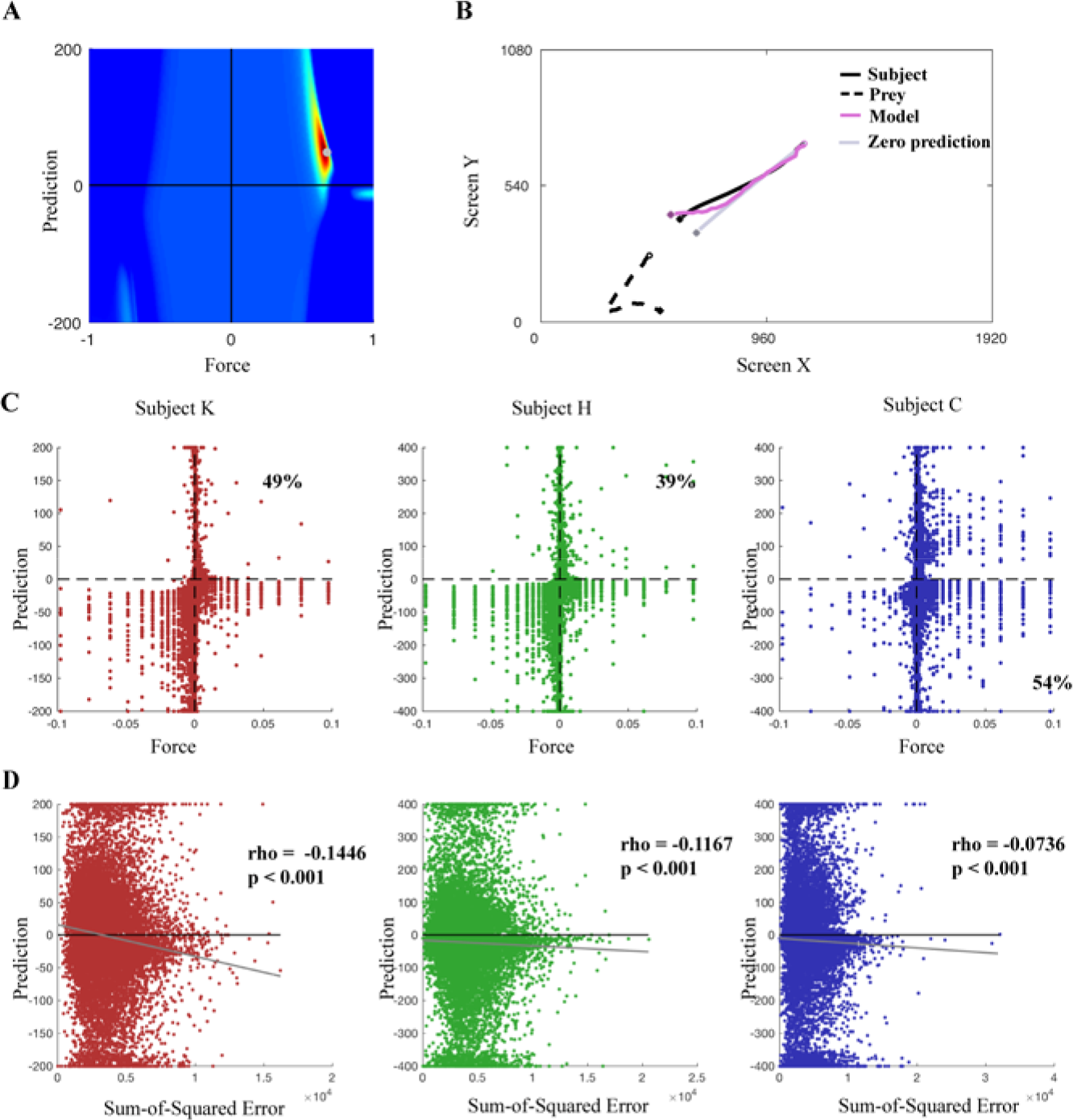
Some trajectories explained with negative prediction. (**A**) Though the representative parameter set appears at positive prediction and positive force (identical figure as PVBP heatmap at figure 2C, across all trajectory heatmap). (**B**) Some trajectories are better explained with negative prediction, as indicated by an example trajectory. The model trajectory in (B) results from PVBP with ‘velocity and acceleration accounted’. (**C**) The fitting of individual trajectory displayed as scatter plot. Percent indicates the most fitting resulted in that quadrant (subject K and H: 1st quadrant with both positive prediction and force; subject C: 4th quadrant with positive force and negative prediction). (**D**) Scatter plot and polynomial fitting (grey solid line) result for cost against prediction. The rho and p-values are obtained from Spearman rank correlation. This provides reason for positive prediction in all subjects: method of heatmap accounts how well each trajectory is fitted.

**Movie 1.**
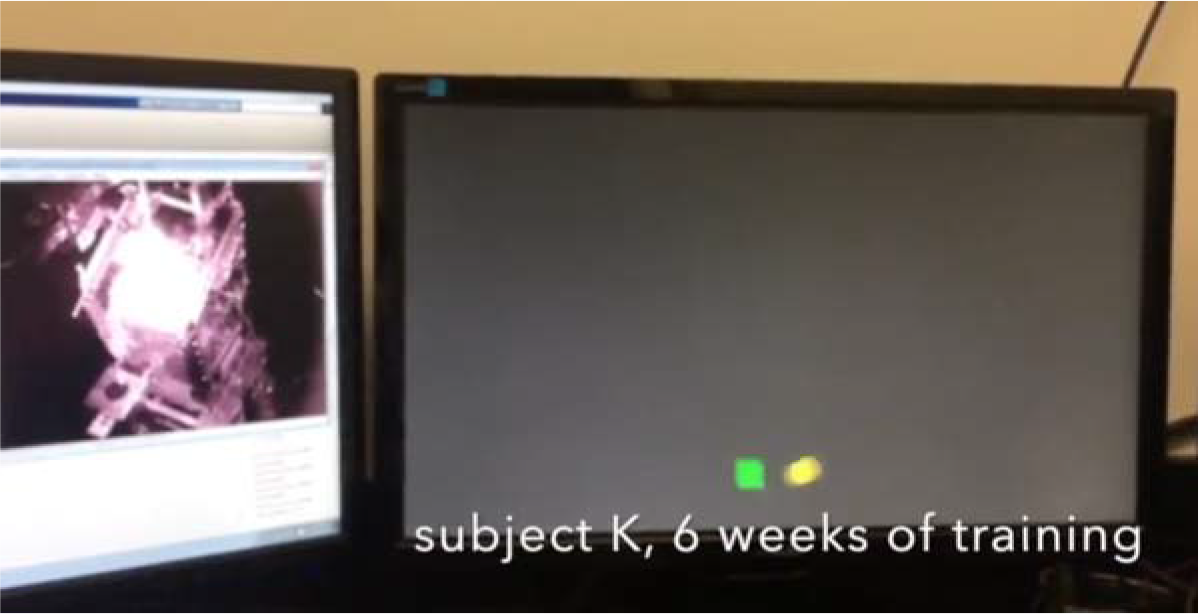
Video of two primates subject playing dynamic pursuit task. Each subject was filmed in different stage of training (subject K: 6 weeks after initial joystick training, subject C: 3 weeks after initial joystick training).

